## Supplementary figures and images for "Small heat shock proteins determine synapse number and neuronal activity during development"

### Supp fig 1

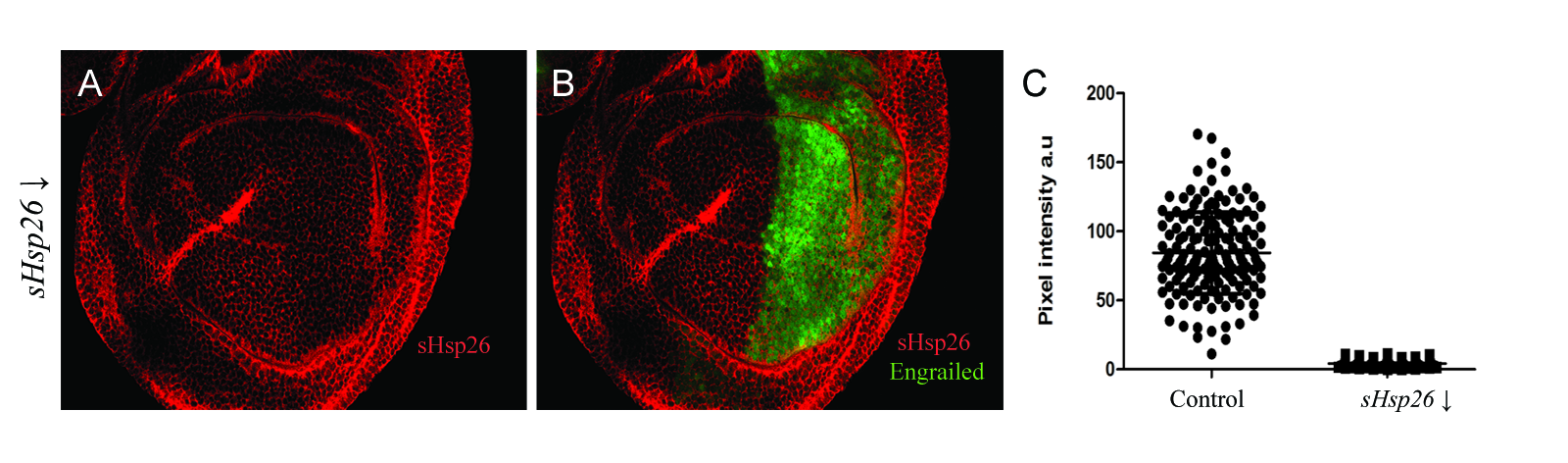
